# Uncoupling of nutrient sensing and cell size control by specific defects in ceramide structure

**DOI:** 10.1101/2025.11.28.691118

**Authors:** José Ignacio Quesada-Márquez, Ana Serrano, María Alcaide-Gavilán, Rafael Lucena

**Affiliations:** Department of Cell Biology. University of Seville. Seville, Spain; Centro Andaluz de Biología del Desarrollo (CABD), CSIC/UPO/JA, Seville, Spain

**Keywords:** Ceramides, cell growth, cell size, yeast, TORC2

## Abstract

Ceramides are essential structural lipids whose chemical diversity arises from variations in acyl-chain length and sphingoid base modifications, yet how these structural features couple metabolic state to growth regulation remains unclear. In *Saccharomyces cerevisiae*, the TORC2–Ypk1/2 signaling axis coordinates plasma membrane homeostasis with cellular growth; however, the precise lipid-derived signals that modulate this pathway remain incompletely characterized. Here, we establish that the structural integrity of very-long-chain fatty acid (VLCFA)–containing ceramides is a critical determinant of nutrient-dependent cell size modulation and TORC2 activity. Disruption of VLCFA elongation elevates Ypk1/2 pT662, consistent with increased TORC2 output, in both nutrient conditions. We further discovered that yeast mutants in VLCFA elongation (*elo3*Δ), fail to modulate their cell size in response to nutrient deprivation. To separate VLCFA elongation from ceramide production, we expressed mammalian ceramide synthases with defined acyl-chain preferences. Ceramides with C_24_–C_26_ chains (CerS3) supported near-normal nutrient-dependent size modulation, whereas production of C_22_–C_24_ ceramides (CerS2) phenocopied elongation mutants, leading to hyperactive TORC2 signaling and defective size regulation. Remarkably, cells producing C_18_ ceramides retained size control despite elevated TORC2 activity, revealing that basal signaling and size regulation can be uncoupled. Furthermore, we identify a distinct role for hydroxylation. While *sur2*Δ mutants exhibited size regulation defects resembling elongation mutants, they retained normal nutrient-responsive TORC2 signaling. Conversely, *scs7*Δ mutants maintained normal size regulation but displayed reduced basal TORC2 activity. This striking uncoupling suggests that while the TORC2 pathway senses acyl chain length, sphingoid base hydroxylation is biophysically required downstream for the execution of cell size remodeling. Our findings demonstrate that distinct structural features of ceramides differentially regulate nutrient sensing, signaling intensity, and the mechanical execution of cell size control.

## 1. Introduction

Cellular growth and size homeostasis depend on the coordinated expansion of the plasma membrane and the synthesis of its constituent lipids. In budding yeast *Saccharomyces cerevisiae*, this coordination is governed by the Target of Rapamycin Complex 2 (TORC2), a conserved protein kinase complex that serves as a master regulator of plasma membrane tension and lipid biosynthesis [1–7]. TORC2 signaling is tightly coupled to sphingolipid metabolism; when sphingolipid levels are low or the membrane is stressed, TORC2 is activated to phosphorylate its downstream effectors, the AGC kinases Ypk1 and Ypk2 [5,8,9]. These kinases, in turn, stimulate ceramide synthase activity and upregulate sphingolipid production, restoring membrane homeostasis [5,8,10].

Sphingolipid metabolism and ceramide synthesis are poorly understood. Sphingolipids, sterols, and glycerophospholipids constitute the primary lipid components found in eukaryotic membranes, with a particular abundance in the plasma membrane. Ceramide is not only a building block of sphingolipids, but also a critical signaling molecule in all eukaryotic organisms that regulates numerous cellular processes such as the cell cycle, cell growth and cell size maintenance [11–15]. For example, ceramides can cluster into ceramide-rich ordered domains to regulate vesicle trafficking [16]. Misregulation of ceramide synthesis is associated with various diseases, including cancer [17,18], cardiovascular disease [19,20] or infections of opportunistic pathogens [21,22].

Sphingolipid biosynthesis begins in the endoplasmic reticulum (ER) with the condensation by a serine palmitoyltransferase (SPT) of serine and palmitoyl-CoA to form 3-ketodihydrosphingosine, which is rapidly reduced to the long-chain base (LCB) dihydrosphingosine (DHS) [23]. Subsequent to the LCB formation, the pathway divides into two distinct arms, the dihydro (non- hydroxylated) and the phyto (hydroxylated) branches. The C4-hydroxylase Sur2 is the enzyme controlling the levels of these two branches [24]. Ceramide synthase subunits Lac1 and Lag1 catalyze N-acylation of DHS to a very long-chain fatty acid (C_26_-VLCFA), producing dihydroceramide (DHCerA) [25–27]. DHCer is hydroxylated at C4 by Sur2 to give phytoceramide (PHCerB). Alternatively, Sur2 can hydroxylate DHS to form PHS, which is then amide linked to the same C_26_-VLCFA to yield phytoceramide [24,28]. Thus, PHCer can be generated through two distinct routes: either via hydroxylation of DHS to PHS followed by N-acylation, or through direct C4-hydroxylation of DHCer by Sur2. DHCer and PHCer can also be hydroxylated at the C2 position of the acyl chain by Scs7 [24,29] to generate DHCerB’ or PHCerC. Finally, the polar head of ceramides can be further modified at the Golgi to generate complex sphingolipids [30]. These include inositol phosphoceramide (IPC), mannosylinositol phosphorylceramide (MIPC) or mannosyldiinositol phosphorylceramide (MIP_2_C).

Of interest, the synthesis of the VLCFAs required for ceramide production comprises a specialized elongation cycle involving Elo1, Elo2, and Elo3. While Elo2 is the rate-limiting enzyme responsible for elongating fatty acids up to C_24_, Elo3 is unique in its ability to catalyze the final extension to C_26_ [31].

Despite the established link between sphingolipid abundance and TORC2 activity, the precise lipid moiety, acyl chain variations or hydroxylation status of ceramides that dictate TORC2 output and cell size regulation remains poorly understood. It remains unclear whether the signaling mechanism relies on sensing bulk lipid levels or recognizing specific structural motifs to couple cell size with nutrient availability. Recent advances have started to delineate the biophysical requirements necessary for this sensing process [5,14,16,32–37]. For example, complete inactivation of ceramide synthases in budding yeast hyperactivates TORC2 signaling despite adding LCB exogenously, which points to ceramide production as a key signaling molecule [5].

In this study, we dissected the functional importance of ceramide structure in regulating cell size and TORC2 signaling in budding yeast. By systematically manipulating ceramide acyl chain length, using both genetic mutants in VLCFA synthesis and by expressing a panel of mammalian ceramide synthases with distinct specificities, we investigated how the cell interprets different ceramide species in the context of nutrient modulation of cell size. Our results suggest that long (C_24_–C_26_) acyl chains with a hydroxylated sphingoid base are sufficient to robustly support basal TORC2 signaling and nutrient-dependent size modulation; however, shorter chains (C_18_–_22_) can permit size remodeling when paired with sphingoid base hydroxylation. We find that ceramides failing to meet this standard are interpreted as a stress signal, leading to growth defects and dysregulated TORC2 signaling. Strikingly, we also uncover that very short-chain ceramides can decouple the regulation of cell size from the control of TORC2’s basal activity, revealing unexpected complexity in how lipid structure is translated into distinct downstream cellular responses.

## 2. Results

### 2.1 Elo3-dependent VLCFA elongation contributes to nutrient-dependent regulation of cell size and is associated with altered TORC2 signaling

Our previous work showed that alterations in ceramide synthesis leads to significant defects in cell size and growth rate [5,14]. This initiates a response to enhance ceramide production through a TORC2-dependent pathway that involves the conserved SGK kinases Ypk1 and Ypk2 and the phosphatase PP2A through its regulatory subunit Rts1 [5,37].

Ceramide biosynthesis requires an amide-linked between a long-chain base (LCB) and a very long-chain fatty acid (VLCFA). A unique and distinguishing feature of ceramides in budding yeast is their C_26_ VLCFA. VLCFAs are produced through a four-step elongation cycle of shorter fatty acids, typically ranging from C_16_ to C_18_ in length. In budding yeast, the Elo1 elongase extends the C_12_-C_16_ fatty acyl-CoAs to the C_16_-C_18_ fatty acids [38]. Elo2 is the rate-limiting enzyme in the elongation of fatty acid to VLCFA [39] using shorter chain fatty acids substrates (C_16_-_18_) and elongates them to acyl-chains maximally containing 24 carbon atoms (C_24_). Elo3, in turn, uses saturated fatty acids of C_18_ as substrates to synthesize VLCFAs in the range of C_20_ to C_26_. Ceramide is formed from the direct condensation of VLCFA with an LCB by the ceramide synthases Lac1 and Lag1 (Figure 1A).

**Figure 1.**
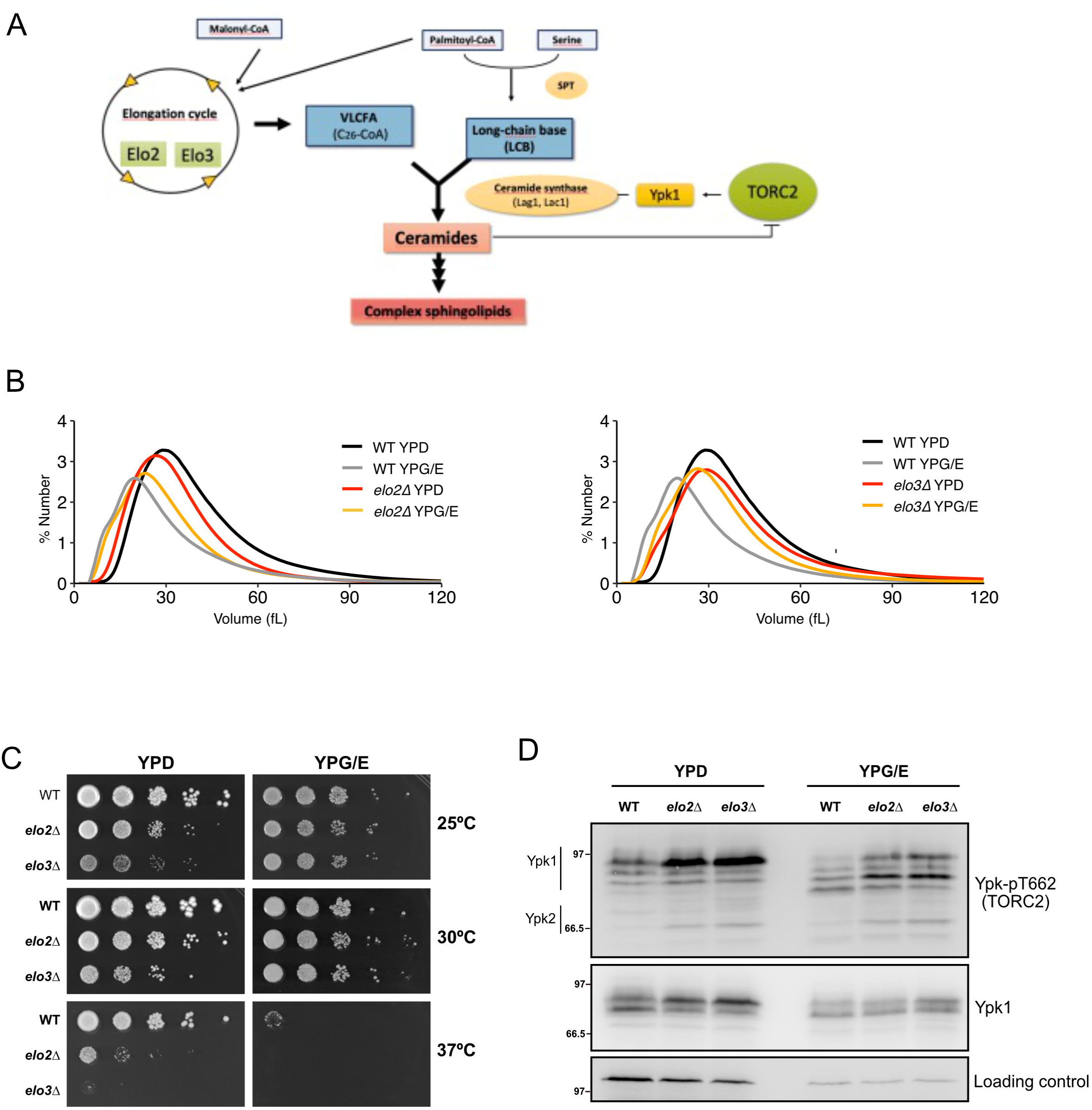
Elo2 and Elo3-dependent VLCFA elongation modulates nutrient-responsive cell size and TORC2 signaling. **(A)** Schematic representation of the fatty acid elongation pathway in *S. cerevisiae*. Elo1 elongates C_12_–C_16_ fatty acids, Elo2 extends C_16_–C_18_ species to C_24_, and Elo3 catalyzes the final step to generate C26 VLCFAs utilized by ceramide synthases Lag1 and Lac1. (**B)** Cells of the indicated genotypes were grown to log phase at 25°C and cell size distributions were determined using a Coulter counter. *elo2*Δ cells exhibit a modest size reduction, whereas *elo3*Δ cells fail to downshift their size in poor carbon conditions. Each plot is the average of 3 biological replicates **(C)** A series of 10-fold dilutions of the indicated strains were grown on YPD or YPG/E plates at 25°C, 30°C and 37°C. **(D)** Cells of the indicated genotypes were grown at 25°C to log phase in YPD or YPG/E medium. Ypk-pT662 and total Ypk1 were assayed by Western blot.

Since *elo1*Δ exhibited an overall fatty acid composition similar to that of a wild type and contains normal amounts of VLCFAs [40], we focused on Elo2 and Elo3 mutants. In rich carbon source (YPD), *elo2*Δ cells showed a small but consistent reduction in cell size compared to wild type, while *elo3*Δ cells were comparable in size to WT. When shifted to a poor carbon source (YPG/E), *elo2*Δ cells adjusted their size to some extent. However, Elo3 was found to be essential for the nutrient modulation of cell size, since *elo3*Δ cells failed to adjust their size (Figure 1B). Interestingly, *elo3*Δ cells exhibited improved colony proliferation at 25°C on poor carbon sources compared to rich (Figure 1C).

We next analyzed TORC2 activity using a phosphospecific antibody (pT662) that recognizes a canonical TORC2-dependent phosphorylation site on its substrates, Ypk1 and Ypk2. The pT662 site is a canonical TORC2 target and widely used as a readout (Niles et al., 2012). Compared to wildtype, pT662 phosphorylation was increased in both *elo2*Δ and *elo3*Δ mutants in rich and poor carbon sources (Figure 1D). This suggests that disrupting VLCFA elongation increases Ypk1/2 pT662, a proxy for TORC2 activity. Overall, these findings indicate that alteration of VLCFAs are sufficient to trigger TORC2 activation, but only the accumulation of C_20_ to C_24_ in *elo3*Δ cells are crucial for the nutrient-dependent regulation of cell size.

### 2.2 Expression of mammalian ceramide synthase homologs differently impacts cell size and TORC2 signaling

To further investigate the role of ceramide chain length we decided to express mammalian ceramide synthases in yeast. In mammalian cells, six ceramide synthases (CerS1-CerS6) have been described so far [41–43]. Unlike the yeast machinery, which produces a limited range of acyl chains, these mammalian isoforms display strict specificity toward fatty acyl-CoAs of defined lengths. Consequently, the expression of CerS homologues in budding yeast resulted in the production of ceramides and sphingolipids with different lengths of the fatty acid chain ([35] and Figure 2A). We focused on ceramide synthases that synthesize major species of fatty acyl-CoA and ceramides ranging from C_18_ to C_26_.

**Figure 2.**
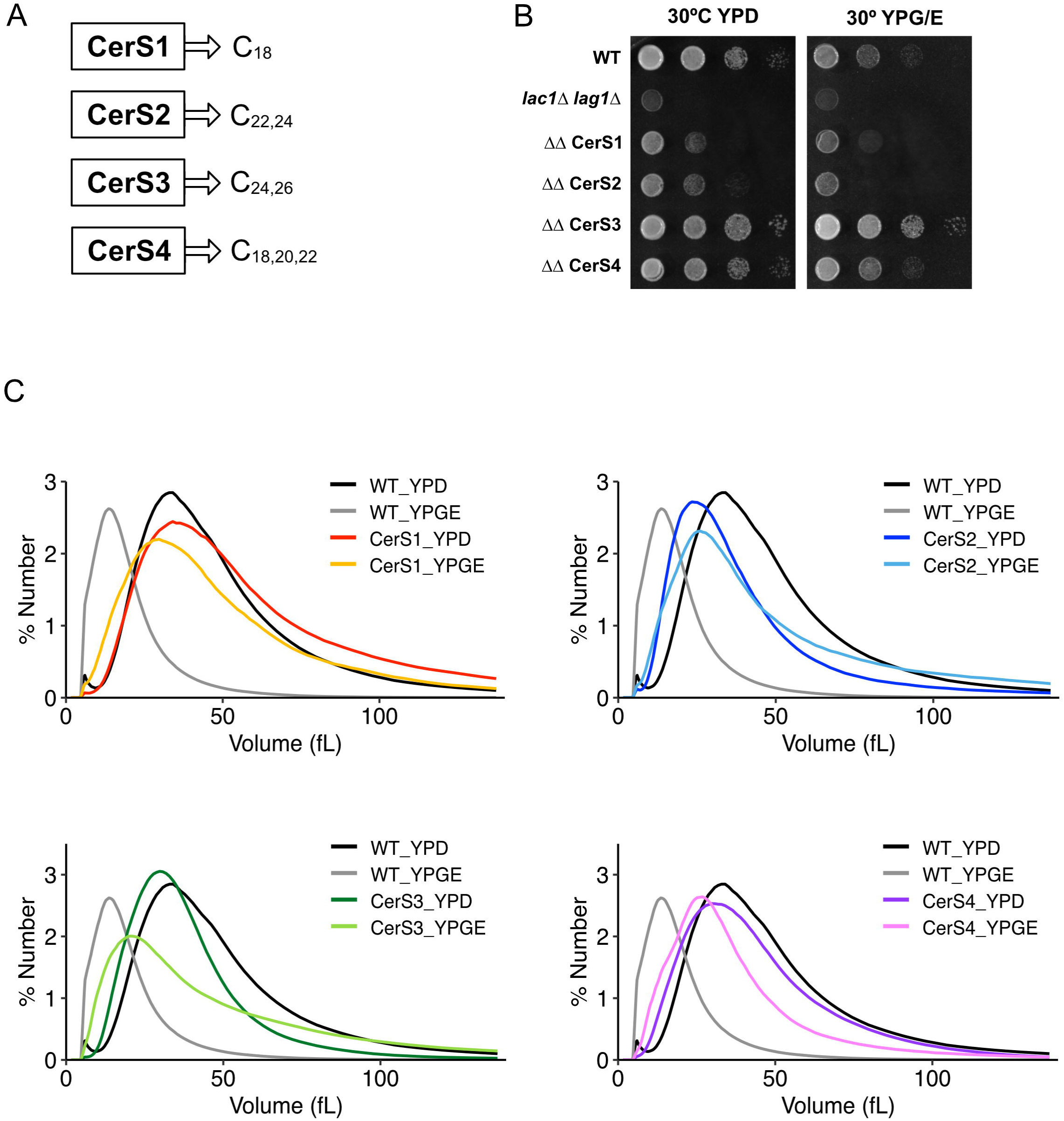
Mammalian CerS isoforms reconstitute distinct ceramide species and produce differential growth and size phenotypes in yeast. **(A)** Fatty acyl-CoA chain-length specificities of CerS1, CerS2, CerS3 and CerS4 according to [44]. **(B)** A series of 10-fold dilutions of the indicated strains were grown on YPD or YPG/E plates at 25°C and 30°C. All CerS isoforms rescue *lac1*Δ *lag1*Δ lethality; CerS1 and CerS2 expression markedly reduces colony growth. **(C)** Cell size distributions of Wild-type and CerS-expressing strains in YPD and YPG/E. CerS2-expressing cells show a severe size reduction and loss of nutrient-dependent size modulation, phenocopying *elo3*Δ. CerS3 and CerS4 produce near-normal size profiles. Each plot is the average of 3 biological replicates.

To confirm that the cellular lipid profile was defined solely by these enzymes, we expressed them in a *lac1*Δ *lag1*Δ double mutant lacking endogenous ceramide synthase activity. All four mammalian CerS rescued the lethality of the *lag1*Δ *lac1*Δ lethality and produced distinct growth and size phenotypes (Figure 2B and [44]), indicating functional output despite species differences. However, strains expressing CerS1 (C_18_ specificity) and CerS2 (C_22_-_24_) grew slower than those expressing CerS3 (C_24_-_26_) and CerS4 (C_18_-_22_). Notably, the CerS1 and CerS2-expressing strains showed a drastic reduction in colony growth both in rich and poor media (Figure 2B).

Analysis of cell volume revealed striking differences. While CerS1 was similar to wildtype in rich media, CerS3 and CerS4 expression resulted in a cell size slightly smaller than wildtype cells. In contrast, expression of CerS2 caused a dramatic decrease in cell size, yielding cells as small as the *lag1*Δ *lac1*Δ mutant (Figure 2C). Furthermore, CerS1, CerS3 and CerS4 reduced their size in response to poor nutrients similarly to *elo2*Δ. Surprisingly, the CerS2 mutant completely failed to modulate its size, similar to our observation with *elo3*Δ mutants (Figure 2C and 1B).

Next, we determined whether these cell size and growth phenotypes correlated with TORC2 activity. The expression of ceramide synthases from mammals was found to have a measurable impact on TORC2 activity to some extent. We found that expression of CerS2, much like the *lag1*Δ *lac1*Δ mutant, caused a significant increase in TORC2-dependent pT662 phosphorylation in both rich and poor media (Figure 3). We also observed a different phosphorylation state of Ypk1 in CerS2 cells in rich media and poor media, similar to the pattern observed in *lac1*Δ *lag1*Δ cells. This may be attributed to the activity of Fpk1/Fpk2 kinases, two redundant kinase paralogues whose roles in regulating Ypk1/2 remain not well understood [45] (Figure 3). These results provide additional evidence that the TORC2 signaling pathway exhibits specificity and adaptation to variations in ceramide acyl chain length. Moreover, it appears that accumulation of ceramides with C_22-24_, as in *elo3*Δ cells, abolished nutrient modulation of cell size.

**Figure 3.**
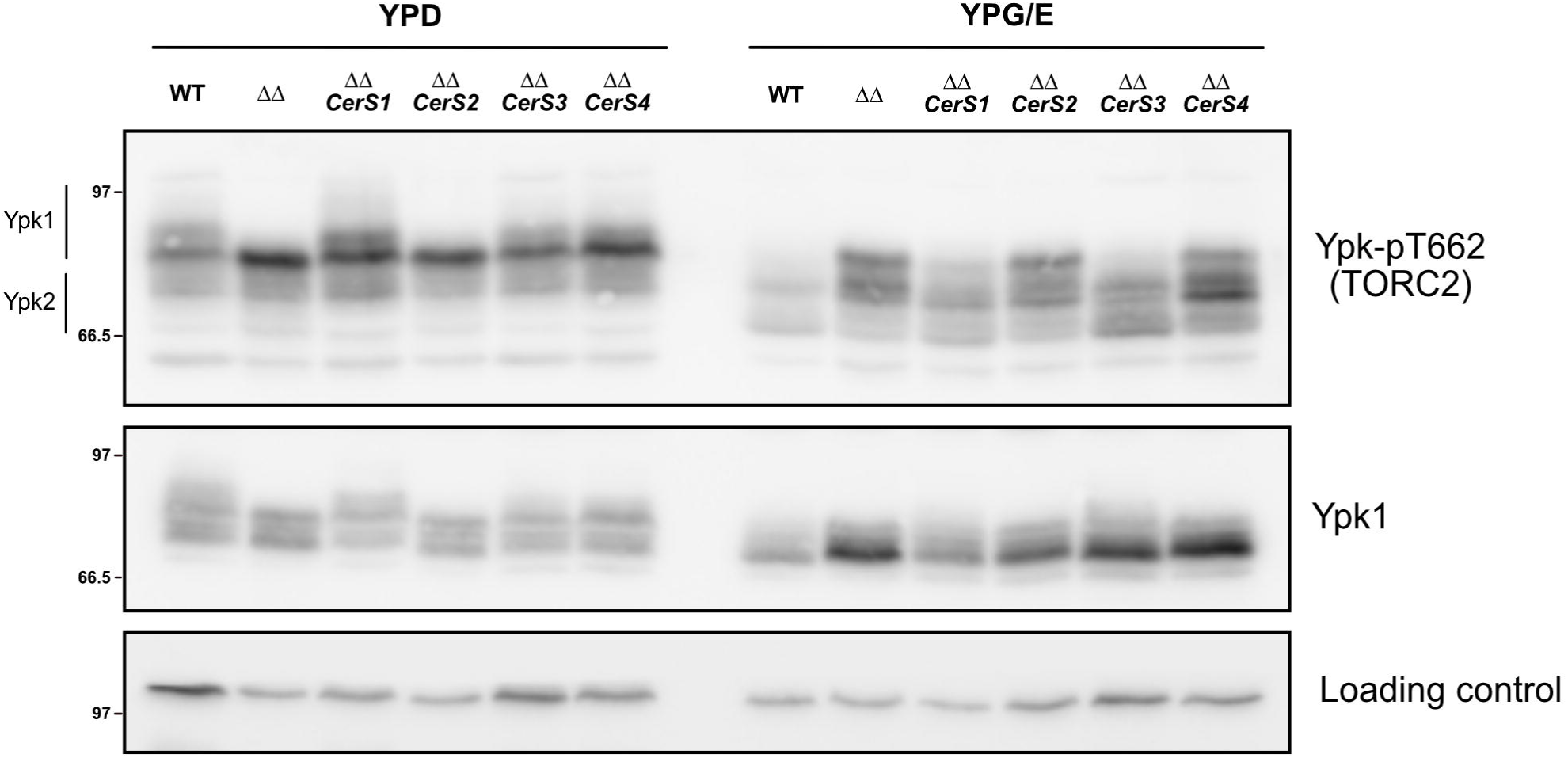
Ceramide acyl-chain length determines TORC2 activation state. Cells of the indicated genotypes were grown at 25°C to log phase in YPD or YPG/E medium. Ypk-pT662 and total Ypk1 were assayed by Western blot.

### 2.3 Short-ceramide mutants decouple cell size modulation from TORC2 signaling

We also explored the evolutionary conservation of this pathway by expressing a codon-optimized cotton Lag1 homolog (GhLag1) in a yeast strain lacking endogenous ceramide synthases. This previously characterized strain produces almost exclusively short-chain C_18_ ceramides [46]. Similar to other short-ceramide mutants, GhLag1 cells showed a consistent decrease in cell size in rich media (Figure 4A). Strikingly, and in contrast to *elo3*Δ and CerS2 mutants, GhLag1 cells were fully capable of adjusting their size in response to a poor carbon source (Figure 4A).

**Figure 4.**
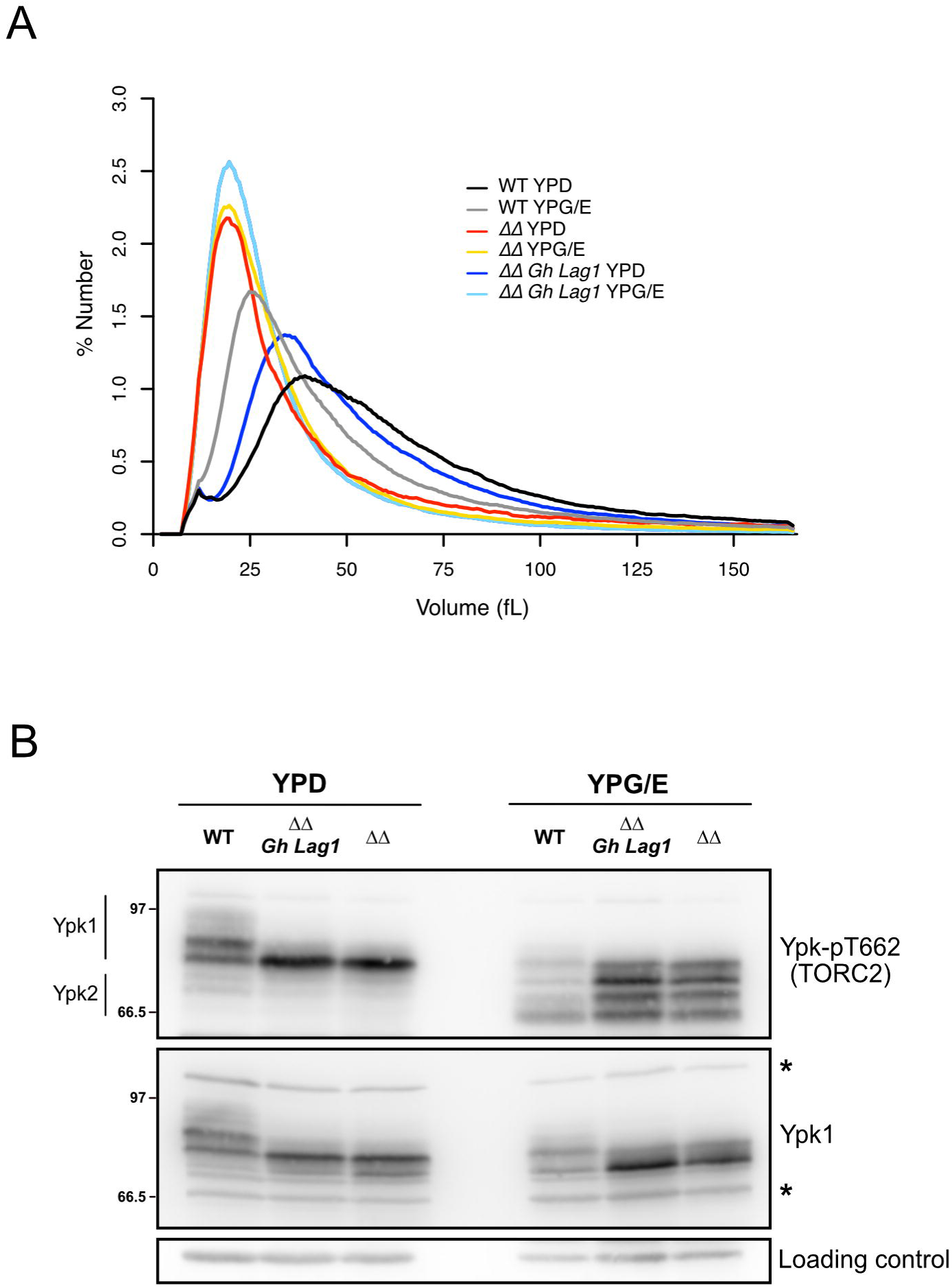
C18 ceramide–producing GhLag1 cells decouple TORC2 signaling from nutrient-dependent cell size control. **(A)** Cells of the indicated genotypes were grown to log phase at 25°C and cell-size distributions were determined using a Coulter counter. Each plot is the average of 3 biological replicates. **(B)** Wild-type, *lac1*Δ *lag1*Δ and GhLag1 cells were grown to early log phase in YPD or YPG/E at 25°C. Ypk-pT662 and total Ypk1 were assayed by Western blot. An asterisk indicates a non-specific band on Ypk1 antibody.

This result was particularly surprising because, despite its normal size modulation, the GhLag1 mutant exhibited highly elevated basal TORC2 signaling (pT662), comparable to the *lag1*Δ *lac1*Δ null mutant, in both rich and poor media (Figure 4B).

We next tested whether TORC2 is modulated in response to rapid changes in carbon source. In wild-type cells, a shift to poor carbon caused rapid loss of TORC2-dependent phosphorylation of Ypk1/2. As cells adapted to the new carbon source, TORC2 activity recovered but remained below levels observed in rich carbon [5]. Thus, we analyzed the acute signaling response to a nutrient shift in GhLag1 cells. In wild-type cells, shifting from rich to poor media causes a rapid and substantial decrease in TORC2 signaling. While GhLag1 cells showed a normal initial drop in pT662, they failed to sustain this response, and TORC2 signaling rapidly recovered to abnormally high levels (Supplemental Figure 1).

Together, these results indicate that short C_18_ ceramides are sufficient to trigger the nutrient-dependent cell size modulation response. However, they are insufficient to properly regulate the basal activity or the acute nutrient-responsive dynamics of the TORC2 signaling network.

### 2.4 Differential roles of sphingoid base and fatty acid hydroxylation in cell size control

To determine if the position of the hydroxyl modification within the ceramide structure dictates it signaling function, we dissected the roles of Sur2 and Scs7 hydroxylases. We compared cell growth, cell size, and TORC2 signaling profiles of single and double deletion mutants in rich (YPD) and poor (YPG/E) carbon sources.

We first assessed cellular growth via serial dilution assays (Figure 5A). The *sur2*Δ and scs7Δ mutants displayed better growth to non-fermentable carbon sources, a phenotype reminiscent of the *elo2*Δ mutant. All mutants also showed a slightly better response to high temperature (37°C). The double mutant *sur2*Δ *scs7*Δ phenocopied the response of the *sur2*Δ single mutant, indicating that the lack of base hydroxylation is the dominant stress determinant.

**Figure 5.**
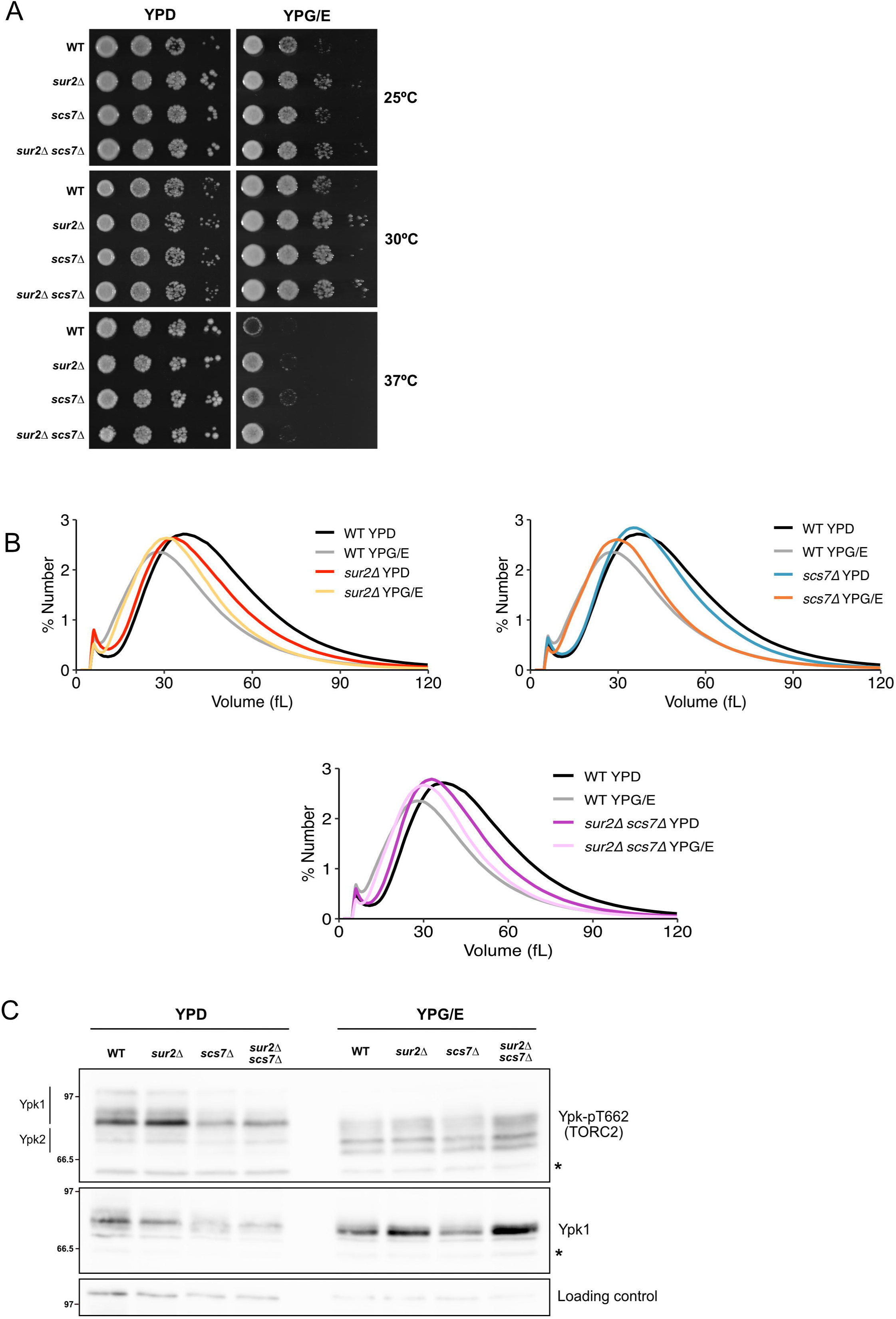
Differential requirements of sphingoid base and fatty acid hydroxylation for cell growth, size homeostasis, and TORC2 signaling. **(A)** Analysis of cell growth. Wild-type (WT), *sur2*Δ, *scs7*Δ, and double mutant *sur2*Δ *scs7*Δ strains were grown to exponential phase, and 10-fold serial dilutions were spotted onto rich (YPD) or poor (YPG/E) media plates. Plates were incubated at the indicated temperatures (25°C, 30°C and 37°C) for 2–3 days. **(B)** Coulter counter cell size analysis. The indicated strains were grown to log phase in YPD or YPG/E. Cell volume distributions (fL) were measured using a Coulter counter as described in methods. Each plot is the average of 3 biological replicates. **(C)** Assessment of TORC2 signaling activity. Whole-cell extracts from strains grown in YPD or YPG/E were analyzed by Western blotting using phosphospecific antibodies against Ypk1-pT662 and total anti-Ypk1 antibodies. An asterisk indicates a non-specific band.

Next, we examined cell size regulation using Coulter counter analysis (Figure 5B). The *scs7*Δ mutant displayed a cell size profile comparable to wild-type. Importantly, these cells retained the ability to modulate their volume, maintaining the ability to perform nutrient modulation of cells size when transferred from YPD to YPG/E. This demonstrates that hydroxylation of the fatty acid is dispensable for nutrient-dependent size remodeling. In contrast, *sur2*Δ cells exhibited a small cell size in rich media and failed to undergo further size reduction in poor media, similar to what we observed in *elo2*Δ cells. Again, the *sur2*Δ *scs7*Δ double mutant showed a size distribution identical to the *sur2*Δ single mutant, confirming that the loss of Sur2 partially prevents size modulation regardless of the Scs7 status.

Finally, we analyzed TORC2 pathway activity (Figure 5C). While *sur2*Δ cells failed to adjust their size, they retained a near normal, nutrient-responsive TORC2 signaling pattern when shifted to YPG/E. Conversely, *scs7*Δ cells, despite their normal size regulation, exhibited reduced basal TORC2 signaling in both rich and poor nutrient conditions. The double mutant mirrored the *sur2*Δ phenotype but with the reduced signaling intensity characteristic of *scs7*Δ.

Taken together, these results suggest a functional hierarchy: Sur2-dependent hydroxylation is biophysically essential for the mechanical execution of cell size control, whereas Scs7-dependent hydroxylation functions primarily to modulate the gain or intensity of the TORC2 signaling output.

## 3. Discussion

The intricate diversity of lipid structures is fundamental to their function, yet how specific structural variations impact complex cellular processes like growth control remains a key question [47–49]. In this study, we provide evidence that the structural integrity of ceramides, specifically their acyl chain length and hydroxylation, is a critical input for the nutrient-sensing TORC2 pathway and the regulation of cell size in *Saccharomyces cerevisiae*. By systematically manipulating acyl chain length, hydroxylation state, and biosynthetic origin of ceramides, we identify the specific lipid features required for nutrient modulation of cell growth and for the proper tuning of TORC2 activity. Our findings support a model where the cell monitors its metabolic state by sensing the presence of mature, correctly structured ceramides and sphingolipids, which in turn dictates decisions on cell growth and proliferation.

Our genetic analysis of VLCFA elongases demonstrates that the final Elo3-dependent elongation step—producing C_26_ acyl chains—is essential for cell size reduction under poor nutrient conditions. Although both *elo2*Δ and *elo3*Δ mutants displayed elevated TORC2 signaling, only *elo3*Δ cells failed to adjust their size upon nutrient limitation. This uncoupling indicates that TORC2 hyperactivation alone cannot compensate for the absence of long-chain ceramides and suggests that specific C_24_–C_26_ acyl chains are biophysically required for membrane remodeling. This result suggests that membranes lacking C_26_ ceramides are perceived by the cell as being under stress, similar to the response observed during the lack of ceramide production [5], membrane tension changes [9,50] and also from VLCFA synthesis [39,51].

The expression of mammalian ceramide synthases reinforced this conclusion [44]. CerS3 (C_24_–C_26_) restored normal cell size and nutrient responsiveness to *lag1*Δ *lac1*Δ cells, whereas CerS2 (C_22_–C_24_) produced extremely small cells incapable of nutrient-dependent size reduction. These observations reveal a structural threshold for function: ceramides shorter than C_24_ are interpreted as a stress signal, whereas C_24_–C_26_ acyl chains are required to maintain proper physiological responses. Because all CerS constructs are expressed from the same TDH3 promoter, differences in phenotypes are unlikely to arise from major differences in expression level. However, we cannot exclude the possibility that variation in protein stability or activity contributes to the observed effects.

A striking finding of our study came from the GhLag1 mutant, which exclusively produces C_18_ ceramides [46]. These cells decoupled the two key phenotypes we measured: nutrient modulation of cell size and TORC2 signaling. While exhibiting constitutively high TORC2 signaling, GhLag1 cells were, surprisingly, still able to reduce their size when shifted to a poor carbon source. This result suggests that the mechanisms governing TORC2 basal activity and nutrient-responsive cell size adjustment can be separated.

The contradictory behavior of the two C_18_-producing enzymes, GhLag1 and CerS1, revealed an unexpected layer of complexity. GhLag1 cells retained the ability to modulate cell size despite having short chains and high TORC2 signaling. In contrast, CerS1 cells failed to modulate size. We propose that this divergence arises from distinct substrate specificities for the LCB. GhLag1 likely utilizes PHS to generate C_18_-PHCer, which satisfy the biophysical requirements for membrane remodeling even if their short length triggers stress signaling. Conversely, mammalian CerS1 may preferentially utilize DHS [52]. This would result in the accumulation of C_18_-DHCer and mechanically impairing size control in response to nutrients. This model suggests that short ceramides can support nutrient modulation of cell size only when they possess the critical C4-hydroxyl group. Notably, CerS1 is structurally and functionally distinctive from the other CerS and is found on an entirely separate branch of the phylogenetic tree [53].

Our analysis of hydroxylation mutants allowed us to distinguish the specific lipid features driving nutrient modulation of cell size from those responsible for adjusting TORC2 signaling output. We found that Scs7-dependent hydroxylation of the fatty acid is dispensable for the fundamental mechanics of cell size regulation. Although *scs7*Δ mutants displayed a reduced TORC2 activity both in rich media and poor media, they retained the ability to modulate their cell size in response to a nutrient shift. This suggests that the hydroxylated phytoceramides generated by Scs7 function primarily as a rheostat to set the amplitude of the TORC2 output, rather than as a binary switch for the response.

In contrast, the loss of Sur2-mediated sphingoid base hydroxylation revealed a fundamental dissociation between signaling and phenotype. *sur2*Δ mutant showed decreased cell size in rich media as previously described [53,54]. It also phenocopied the defects of elongation mutants yet retained a normal TORC2-signaling in response to nutrient shift. We conclude that the defect observed in *sur2*Δ cells may represent a failure of biophysical competence rather than sensory perception; the signaling machinery detects the nutrient shift, but the membrane architecture is unable to execute the required morphological change.

Altogether, our findings support a model in which yeast cells monitor the structural integrity of ceramides, rather than their bulk levels, to coordinate membrane composition with growth decisions. Mature C_26_-PHCer, defined by a long acyl chain and a hydroxylated LCB, act as a molecular signal of membrane health. Deviations from this molecular signature—whether due to shortened acyl chains, loss of LCB hydroxylation, or incompatibility of heterologous ceramide synthases—activate TORC2 and impair cell size control to some extent.

This structural model explains how TORC2 converts subtle lipid structural features into distinct physiological responses and raises new questions about the molecular sensors responsible for detecting ceramide structure. Our work establishes that ceramide structure is a critical determinant of TORC2 signaling and nutrient modulation of cell size control in budding yeast. By identifying the specific structural motifs required for proper function, we reveal how membrane lipid architecture is translated into actionable growth signals. Future studies aimed at identifying the molecular machinery that senses ceramide structure and linking these lipids to TORC2 regulation will provide fundamental insight into conserved principles of lipid-based signaling across eukaryotes.

## 4. Materials and methods

### 4.1 Yeast strains and culture conditions

All strains are in the W303 background (leu2-3,112 ura3-1 can1-100 ade2-1 his3-11,15 trp1-1 GAL + ssd1-d2). Table 1 shows additional genetic features and backgrounds. One-step PCR-based gene replacement was used for making deletions at the endogenous locus [55,56]. Cells were grown in YP medium (1% yeast extract, 2% peptone, 40 mg/liter adenine) supplemented with 2% dextrose (YPD) or 2% glycerol/ethanol (YPG/E). To analyze cells shifted from rich to poor nutrients (Supplementary Figure 1), cultures were grown in YPD medium overnight at 25°C to an OD_600_ of less than 0.8. After adjusting optical densities to normalize protein loading, cells were washed three times with a large volume of YPG/E medium and then incubated at 30°C in YPG/E for the time course. 1.6 ml samples were collected at each time point.

**Table 1.** Strains used in this study.

| Strain | MAT | Genotype | Source |
| --- | --- | --- | --- |
| RL1 | a | *hur1* | Douglas Kellogg |
| RL327 | a | *elo2Δ::KanMX* | This study |
| RL337 | a | *elo3Δ::KanMX* | This study |
| AL378 | n | *lag1Δ::HIS3 lac1Δ::ADE2* | Howard Riezman (RH7165) |
| AL379 | n | RH7165 TDH3<sub>pu</sub>-CERS1-CYC1 term TRP1 | Howard Riezman (RH7911) |
| AL380 | n | RH7165 TDH3<sub>pu</sub>-CERS2-CYC1 term TRP1 | Howard Riezman (RH7912) |
| AL381 | n | RH7165 TDH3<sub>pu</sub>-CERS3-CYC1 term TRP1 | Howard Riezman (RH7913) |
| AL382 | n | RH7165 TDH3<sub>pu</sub>-CERS4-CYC1 term TRP1 | Howard Riezman (RH7914) |
| RL190 | a | *lag1Δ::HIS3 lac1Δ::ADE2 TDH3::GhLAG1::TRP1* | MMY1678 |
| AL402 | a | *sur2Δ::HIS* | This study |
| AL445 | a | *scs7Δ::HIS* | This study |
| AL435 | a | *sur2Δ::HIS scs7Δ::KanMX* | This study |

Howard Riezman background: *his3-11,15, leu2, trp1, ura3, ade2, ben1*
MMY1678 background: *his4, leu2, ura3*

### 4.2 Analysis of cell size and cell proliferation assays

Cell cultures were grown overnight to early log phase at 25°C. A 900 µl sample of each culture was fixed with 100 µl of 37% formaldehyde for 30 min and then washed twice with PBS containing 0.04% sodium azide and 0.02% Tween-20. Cell size was measured using a Coulter Counter Z3 (Channelizer Z3, Beckman Coulter) as previously described [5]. In brief, cells were diluted into 10 ml diluent (Isoton II; Beckman Coulter) and sonicated for 3 s before cell sizing. Each plot is the average of three independent biological replicates in which three independent technical replicates were averaged. To assay the rate of cell proliferation on plates, cells were grown overnight in the indicated medium at 25°C and adjusted to an OD_600_ of 0.5. Tenfold serial dilutions were spotted onto YPD or YPG/E and incubated at 25°C, 30°C or 37°C for 3 days.

### 4.3 Western blotting

For Western blotting, cells growing in early log phase were grown overnight at 25°C to an OD_600_ of 0.6 as described in [57]. After adjusting optical densities to normalize protein loading, 1.6 ml samples were collected and centrifuged at 20,000 g for 30 sec. The supernatant was removed and 250 µl of glass beads were added before freezing in liquid nitrogen. We collected 1.6 ml samples at each time point. Cells were lysed into 140 µl of sample buffer (65 mM Tris HCl, pH 6.8, 3% SDS, 10% glycerol, 50 mM sodium fluoride, 100 mM β-glycerophosphate, 5% 2-mercaptoethanol, and bromophenol blue). PMSF was added to the sample buffer to 2 mM immediately before use. Cells were lysed in a Mini-beadbeater 16 (BioSpec) at top speed for 2 min. The samples were removed and centrifuged for 15 sec at 20,000 g in a microfuge and placed in boiling water for 5 min. After boiling, the samples were centrifuged for 5 min at 20,000 g and loaded onto an SDS polyacrylamide gel. Samples were analyzed by Western blotting as previously described [58]. SDS-PAGE gels were run at a constant current of 20 mA and electrophoresis was performed on gels containing 10% polyacrylamide and 0.13% bis-acrylamide. Proteins were transferred to nitrocellulose using a Trans-Blot Turbo system (Bio-Rad). Blots were probed with primary antibody overnight at 4°C. Rabbit anti–phospho-T662 antibody (a kindly gift from Ted Powers, University of California, Davis) was used to detect TORC2-dependent phosphorylation of Ypk1/2 at a dilution of 1:20,000 in TBST (10 mM Tris-Cl, pH 7.5, 100 mM NaCl, and 0.1% Tween 20) containing 3% milk. Total Ypk1 was detected using anti-Ypk1 antibody (a kindly gift from Douglas Kellogg, [59]) at a dilution of 1:10,000. All blots were probed with an Anti-Rabbit IgG Peroxidase AffiniPure Goat secondary antibody (Jackson ImmunoResearch, 111-035-003) for 45–90 min at room temperature.

## Author contributions

Conceptualization: RL; Investigation: JIQM, AS, MAG, RL; Methodology: MAG, RL; Validation: JIQM, RL; Visualization: MAG; Funding acquisition: MAG, RL; Project administration: RL; Resources: MAG, RL; Supervision: RL; Writing- original draft: RL; Writing- review and editing: JIQM, AS, MAG, RL.

## Acknowledgements

We thank members of the laboratory for advice and support. We thank Ted Powers (University of California, Davis) for the Ypk-pT662 phosphospecific antibody, Manuel Muñiz (University of Seville) for strains, reagents and material and Howard Riezman (University of Geneva) for strains. We thank Sebastian Chávez and María Cruz Muñoz-Centeno (University of Seville) for using the Coulter Counter Z3 and Derek McCusker (University of Bordeaux) for materials. We are deeply grateful to Douglas Kellogg (University of California, Santa Cruz) for Ypk1 antibody, reagents, strains and his support during a short stay in his laboratory.

---

**Supplemental Figure 1. Defective TORC2 dynamics in GhLag1 cells following acute nutrient downshift.**

Time course of Ypk1/2 pT662 phosphorylation in WT and GhLag1 cells after a shift from YPD to YPG/E. WT cells show a rapid decrease followed by partial recovery. GhLag1 cells exhibit an initial decrease but rapidly return to elevated TORC2 activity, indicating impaired regulation of acute nutrient-responsive TORC2 dynamics.

## REFERENCES

[1] A. Emmerstorfer-Augustin, J. Thorner, Regulation of TORC2 Function and Localization in Yeast, Annu Rev Cell Dev Biol 39 (2023). 10.1146/annurev-cellbio-011723-030346.

[2] M. Riggi, B. Kusmider, R. Loewith, The flipside of the TOR coin – TORC2 and plasma membrane homeostasis at a glance, J Cell Sci 133 (2020) jcs242040. 10.1242/jcs.242040.

[3] F. Roelants, K. Leskoske, M.N. Martinez Marshall, M. Locke, J. Thorner, The TORC2[Dependent Signaling Network in the Yeast Saccharomyces cerevisiae, Biomolecules 7 (2017) 66. 10.3390/biom7030066.

[4] J. Thorner, TOR complex 2 is a master regulator of plasma membrane homeostasis, Biochemical Journal 479 (2022) 1917–1940. 10.1042/BCJ20220388.

[5] R. Lucena, M. Alcaide-Gavilán, K. Schubert, M. He, M.G. Domnauer, C. Marquer, C. Klose, M.A. Surma, D.R. Kellogg, Cell Size and Growth Rate Are Modulated by TORC2-Dependent Signals, Current Biology 28 (2018) 196–210.e4. 10.1016/j.cub.2017.11.069.

[6] T. Powers, S. Aronova, B. Niles, TORC2 and Sphingolipid Biosynthesis and Signaling, in: Enzymes, 2010: pp. 177–197. 10.1016/S1874-6047(10)27010-5.

[7] S. Aronova, K. Wedaman, P.A. Aronov, K. Fontes, K. Ramos, B.D. Hammock, T. Powers, Regulation of Ceramide Biosynthesis by TOR Complex 2, Cell Metab 7 (2008) 148–158. 10.1016/j.cmet.2007.11.015.

[8] F.M. Roelants, D.K. Breslow, a. Muir, J.S. Weissman, J. Thorner, Protein kinase Ypk1 phosphorylates regulatory proteins Orm1 and Orm2 to control sphingolipid homeostasis in Saccharomyces cerevisiae, Proceedings of the National Academy of Sciences 108 (2011) 19222–19227. 10.1073/pnas.1116948108.

[9] D. Berchtold, M. Piccolis, N. Chiaruttini, I. Riezman, H. Riezman, A. Roux, T.C. Walther, R. Loewith, Plasma membrane stress induces relocalization of Slm proteins and activation of TORC2 to promote sphingolipid synthesis., Nat Cell Biol 14 (2012) 542–7. 10.1038/ncb2480.

[10] A. Muir, S. Ramachandran, F.M. Roelants, G. Timmons, J. Thorner, TORC2-dependent protein kinase Ypk1 phosphorylates ceramide synthase to stimulate synthesis of complex sphingolipids., Elife 3 (2014) 1–34. 10.7554/eLife.03779.

[11] Y.A. Hannun, L.M. Obeid, Sphingolipids and their metabolism in physiology and disease, Nat Rev Mol Cell Biol 19 (2018) 175–191. 10.1038/nrm.2017.107.

[12] C. Chalfant, M. Del Poeta, eds., Sphingolipids as Signaling and Regulatory Molecules, Springer New York, New York, NY, 2010. 10.1007/978-1-4419-6741-1.

[13] S.A. Summers, B. Chaurasia, W.L. Holland, Metabolic Messengers: ceramides, Nat Metab 1 (2019) 1051–1058. 10.1038/s42255-019-0134-8.

[14] I. Flor-Parra, S. Sabido-Bozo, A. Ikeda, K. Hanaoka, A. Aguilera-Romero, K. Funato, M. Muñiz, R. Lucena, The ceramide synthase subunit lac1 regulates cell growth and size in fission yeast, Int J Mol Sci 23 (2022) 1–14. 10.3390/ijms23010303.

[15] A. Alonso, F.M. Goñi, The Physical Properties of Ceramides in Membranes, Annu Rev Biophys 47 (2018) 633–654. 10.1146/annurev-biophys-070317-033309.

[16] A. Aguilera-Romero, R. Lucena, S. Sabido-Bozo, M. Muñiz, Impact of sphingolipids on protein membrane trafficking, Biochimica et Biophysica Acta (BBA) - Molecular and Cell Biology of Lipids 1868 (2023) 159334. 10.1016/j.bbalip.2023.159334.

[17] S.A. Morad, M.C. Cabot, Ceramide-orchestrated signalling in cancer cells, Nat Rev Cancer 13 (2013) 51–65. 10.1038/nrc3398.

[18] J. Alizadeh, S.C. da Silva Rosa, X. Weng, J. Jacobs, S. Lorzadeh, A. Ravandi, R. Vitorino, S. Pecic, A. Zivkovic, H. Stark, S. Shojaei, S. Ghavami, Ceramides and ceramide synthases in cancer: Focus on apoptosis and autophagy, Eur J Cell Biol 102 (2023) 151337. 10.1016/j.ejcb.2023.151337.

[19] C.D. Green, M. Maceyka, L.A. Cowart, S. Spiegel, Sphingolipids in metabolic disease: The good, the bad, and the unknown, Cell Metab 33 (2021) 1293–1306. 10.1016/j.cmet.2021.06.006.

[20] B. Chaurasia, S.A. Summers, Ceramides in Metabolism: Key Lipotoxic Players, Annu Rev Physiol 83 (2021) 303–330. 10.1146/annurev-physiol-031620-093815.

[21] M. Kumar, A. Singh, S. Kumari, P. Kumar, M. Wasi, A.K. Mondal, S.M. Rudramurthy, A. Chakrabarti, N.A. Gaur, N.A.R. Gow, R. Prasad, Sphingolipidomics of drug resistant Candida auris clinical isolates reveal distinct sphingolipid species signatures, Biochim Biophys Acta Mol Cell Biol Lipids 1866 (2021) 158815. 10.1016/j.bbalip.2020.158815.

[22] A.K. Urbanek, J. Muraszko, D. Derkacz, M. Łukaszewicz, P. Bernat, A. Krasowska, The Role of Ergosterol and Sphingolipids in the Localization and Activity of Candida albicans’ Multidrug Transporter Cdr1p and Plasma Membrane ATPase Pma1p, Int J Mol Sci 23 (2022). 10.3390/ijms23179975.

[23] T. Beeler, D. Bacikova, K. Gable, L. Hopkins, C. Johnson, H. Slife, T. Dunn, The Saccharomyces cerevisiae TSC10/YBR265W gene encoding 3-ketosphinganine reductase is identified in a screen for temperature-sensitive suppressors of the Ca2+-sensitive csg2Δ mutant, Journal of Biological Chemistry 273 (1998) 30688–30694. 10.1074/jbc.273.46.30688.

[24] D. Haak, K. Gable, T. Beeler, T. Dunn, Hydroxylation of Saccharomyces cerevisiae ceramides requires Sur2p and Scs7p, Journal of Biological Chemistry 272 (1997) 29704–29710. 10.1074/jbc.272.47.29704.

[25] I. Guillas, P.A. Kirchman, R. Chuard, M. Pfefferli, J.C. Jiang, S.M. Jazwinski, A. Conzelmann, C26-CoA-dependent ceramide synthesis of Saccharomyces cerevisiae is operated by Lag1p and Lac1p, EMBO Journal 20 (2001) 2655–2665. 10.1093/emboj/20.11.2655.

[26] S. Schorling, B. Vallée, W.P. Barz, H. Riezman, D. Oesterhelt, Lag1p and Lac1p Are Essential for the Acyl-CoA–dependent Ceramide Synthase Reaction in Saccharomyces cerevisae, Mol Biol Cell 12 (2001) 3417–3427. 10.1091/mbc.12.11.3417.

[27] B. Vallée, H. Riezman, Lip1p: a novel subunit of acyl-CoA ceramide synthase., EMBO J 24 (2005) 730–41. 10.1038/sj.emboj.7600562.

[28] M.M. Grilley, S.D. Stock, R.C. Dickson, R.L. Lester, J.Y. Takemoto, Syringomycin action gene SYR2 is essential for sphingolipid 4- hydroxylation in Saccharomyces cerevisiae, Journal of Biological Chemistry 273 (1998) 11062–11068. 10.1074/jbc.273.18.11062.

[29] T.M. Dunn, D. Haak, E. Monaghan, T.J. Beeler, Synthesis of monohydroxylated inositolphosphorylceramide (IPC-C) in Saccharomyces cerevisiae requires Scs7p, a protein with both a cytochrome b5-like domain and a hydroxylase/desaturase domain, Yeast 14 (1998) 311–321. 10.1002/(SICI)1097-0061(19980315)14:4<311::AID-YEA220>3.0.CO;2-B.

[30] K. Funato, H. Riezman, Vesicular and nonvesicular transport of ceramide from ER to the Golgi apparatus in yeast, Journal of Cell Biology 155 (2001) 949–959. 10.1083/jcb.200105033.

[31] C.S. Oh, D.A. Toke, S. Mandala, C.E. Martin, ELO2 and ELO3, homologues of the Saccharomyces cerevisiae ELO1 gene, function in fatty acid elongation and are required for sphingolipid formation, Journal of Biological Chemistry 272 (1997) 17376–17384. 10.1074/jbc.272.28.17376.

[32] T. Harayama, H. Riezman, Understanding the diversity of membrane lipid composition, Nat Rev Mol Cell Biol 19 (2018) 281–296. 10.1038/nrm.2017.138.

[33] M.G. Tettamanti, P. Nowak, B. Kusmider, J.M. Kefauver, V. Mercier, A. Roux, R. Loewith, TORC2-regulated sterol redistribution mediates recovery from membrane perturbation by small amphipathic molecules., (2024). 10.1101/2024.10.18.618785.

[34] D.R. Kellogg, P.A. Levin, Nutrient availability as an arbiter of cell size, Trends Cell Biol 32 (2022) 908–919. 10.1016/j.tcb.2022.06.008.

[35] A. Aguilera-Romero, C. Gehin, H. Riezman, Sphingolipid homeostasis in the web of metabolic routes, Biochim Biophys Acta Mol Cell Biol Lipids 1841 (2014) 647–656. 10.1016/j.bbalip.2013.10.014.

[36] S. Rodriguez-Gallardo, K. Kurokawa, S. Sabido-Bozo, A. Cortes-Gomez, A. Ikeda, V. Zoni, A. Aguilera-Romero, A.M. Perez-Linero, S. Lopez, M. Waga, M. Araki, M. Nakano, H. Riezman, K. Funato, S. Vanni, A. Nakano, M. Muñiz, Ceramide chain length-dependent protein sorting into selective endoplasmic reticulum exit sites, Sci Adv 6 (2020). 10.1126/sciadv.aba8237.

[37] M. Alcaide-Gavilán, R. Lucena, K.A. Schubert, K.L. Artiles, J. Zapata, D.R. Kellogg, Modulation of TORC2 Signaling by a Conserved Lkb1 Signaling Axis in Budding Yeast, Genetics 210 (2018) 155–170. 10.1534/genetics.118.301296.

[38] C.S. Oh, D.A. Toke, S. Mandala, C.E. Martin, ELO2 and ELO3, homologues of the Saccharomyces cerevisiae ELO1 gene, function in fatty acid elongation and are required for sphingolipid formation, Journal of Biological Chemistry 272 (1997) 17376–17384. 10.1074/jbc.272.28.17376.

[39] D.K. Olson, F. Fröhlich, R. V. Farese, T.C. Walther, Taming the sphinx: Mechanisms of cellular sphingolipid homeostasis, Biochim Biophys Acta Mol Cell Biol Lipids (2015). 10.1016/j.bbalip.2015.12.021.

[40] F. Dittrich, D. Zajonc, K. Hühne, U. Hoja, A. Ekici, E. Greiner, H. Klein, J. Hofmann, J.J. Bessoule, P. Sperling, E. Schweizer, Fatty acid elongation in yeast - Biochemical characteristics of the enzyme system and isolation of elongation-defective mutants, Eur J Biochem 252 (1998) 477–485. 10.1046/j.1432-1327.1998.2520477.x.

[41] K. Venkataraman, A.H. Futerman, Ceramide as a second messenger: Sticky solutions to sticky problems, Trends Cell Biol 10 (2000) 408–412. 10.1016/S0962-8924(00)01830-4.

[42] I. Guillas, J.C. Jiang, C. Vionnet, C. Roubaty, D. Uldry, R. Chuard, J. Wang, S.M. Jazwinski, A. Conzelmann, Human Homologues of LAG1 Reconstitute Acyl-CoA-dependent Ceramide Synthesis in Yeast, Journal of Biological Chemistry 278 (2003) 37083–37091. 10.1074/jbc.M307554200.

[43] M. Levy, A.H. Futerman, Mammalian ceramide synthases, IUBMB Life 62 (2010) 347–356. 10.1002/iub.319.

[44] J.T. Hannich, A.G. Haribowo, S. Gentina, M. Paillard, L. Gomez, B. Pillot, H. Thibault, D. Abegg, N. Guex, A. Zumbuehl, A. Adibekian, M. Ovize, J.C. Martinou, H. Riezman, 1-Deoxydihydroceramide causes anoxic death by impairing chaperonin-mediated protein folding, Nat Metab 1 (2019) 996–1008. 10.1038/s42255-019-0123-y.

[45] F.M. Roelants, A.G. Baltz, A.E. Trott, S. Fereres, J. Thorner, A protein kinase network regulates the function of aminophospholipid flippases., Proc Natl Acad Sci U S A 107 (2010) 34–39. 10.1073/pnas.0912497106.

[46] S. Epstein, G.A. Castillon, Y. Qin, H. Riezman, An essential function of sphingolipids in yeast cell division, Mol Microbiol 84 (2012) 1018–1032. 10.1111/j.1365-2958.2012.08087.x.

[47] M. Tani, Biological Importance of Complex Sphingolipids and Their Structural Diversity in Budding Yeast Saccharomyces cerevisiae, Int J Mol Sci 25 (2024). 10.3390/ijms252212422.

[48] T. Harayama, B. Antonny, Beyond Fluidity: The Role of Lipid Unsaturation in Membrane Function, Cold Spring Harb Perspect Biol 15 (2023). 10.1101/cshperspect.a041409.

[49] A.H. Futerman, Y.A. Hannun, The complex life of simple sphingolipids, EMBO Rep 5 (2004) 777–782. 10.1038/sj.embor.7400208.

[50] M. Riggi, K. Niewola-Staszkowska, N. Chiaruttini, A. Colom, B. Kusmider, V. Mercier, S. Soleimanpour, M. Stahl, S. Matile, A. Roux, R. Loewith, Decrease in plasma membrane tension triggers PtdIns(4,5)P2 phase separation to inactivate TORC2, Nat Cell Biol 20 (2018) 1043–1051. 10.1038/s41556-018-0150-z.

[51] D.K. Olson, F. Fröhlich, R. Christiano, H.K. Hannibal-Bach, C.S. Ejsing, T.C. Walther, Rom2-dependent phosphorylation of Elo2 controls the abundance of very long-chain fatty acids, Journal of Biological Chemistry 290 (2015) 4238–4247. 10.1074/jbc.M114.629279.

[52] T.D. Mullen, Y.A. Hannun, L.M. Obeid, Ceramide synthases at the centre of sphingolipid metabolism and biology, Biochemical Journal 441 (2012) 789–802. 10.1042/BJ20111626.

[53] Y. Pewzner-Jung, S. Ben-Dor, A.H. Futerman, When do Lasses (longevity assurance genes) become CerS (ceramide synthases)? Insights into the regulation of ceramide synthesis, Journal of Biological Chemistry 281 (2006) 25001–25005. 10.1074/jbc.R600010200.

[54] Z. Deng, Q. Wang, R. Ding, W. Nie, X. Chen, Y. Chen, Y. Wang, J. Duan, Z. Hu, Loss of SUR2 alters the composition of ceramides and shortens chronological lifespan of Saccharomyces cerevisiae, Biochim Biophys Acta Mol Cell Biol Lipids 1870 (2025). 10.1016/j.bbalip.2024.159591.

[55] M.S. Longtine, A. McKenzie, D.J. Demarini, N.G. Shah, A. Wach, A. Brachat, P. Philippsen, J.R. Pringle, Additional modules for versatile and economical PCR-based gene deletion and modification in Saccharomyces cerevisiae, Yeast 14 (1998) 953–961. http://doi.wiley.com/10.1002/(SICI)1097-0061(199807)14:10<953::AID-YEA293>3.0.CO;2-U.

[56] C. Janke, M.M. Magiera, N. Rathfelder, C. Taxis, S. Reber, H. Maekawa, A. Moreno-Borchart, G. Doenges, E. Schwob, E. Schiebel, M. Knop, A versatile toolbox for PCR-based tagging of yeast genes: New fluorescent proteins, more markers and promoter substitution cassettes, Yeast 21 (2004) 947–962. 10.1002/yea.1142.

[57] R. Lucena, A. Jasani, S. Anastasia, D. Kellogg, M. Alcaide-Gavilan, Casein kinase 1 controls components of a TORC2 signaling network in budding yeast, J Cell Sci 137 (2024) 2024.01.30.578072. 10.1242/jcs.262036.

[58] S.L. Harvey, G. Enciso, N. Dephoure, S.P. Gygi, J. Gunawardena, D.R. Kellogg, A phosphatase threshold sets the level of Cdk1 activity in early mitosis in budding yeast., Mol Biol Cell 22 (2011) 3595–3608. 10.1091/mbc.E11-04-0340.

[59] M. Alcaide-Gavilán, R. Lucena, S. Banuelos, D.R. Kellogg, Conserved Ark1-related kinases function in a TORC2 signaling network, Mol Biol Cell 31 (2020) 2057–2069. 10.1091/mbc.E19-12-0685.

