## Supplementary figures and images for "Uncoupling of nutrient sensing and cell size control by specific defects in ceramide structure"

### Supplemental Figure 1

#### Supplementary Figure 1

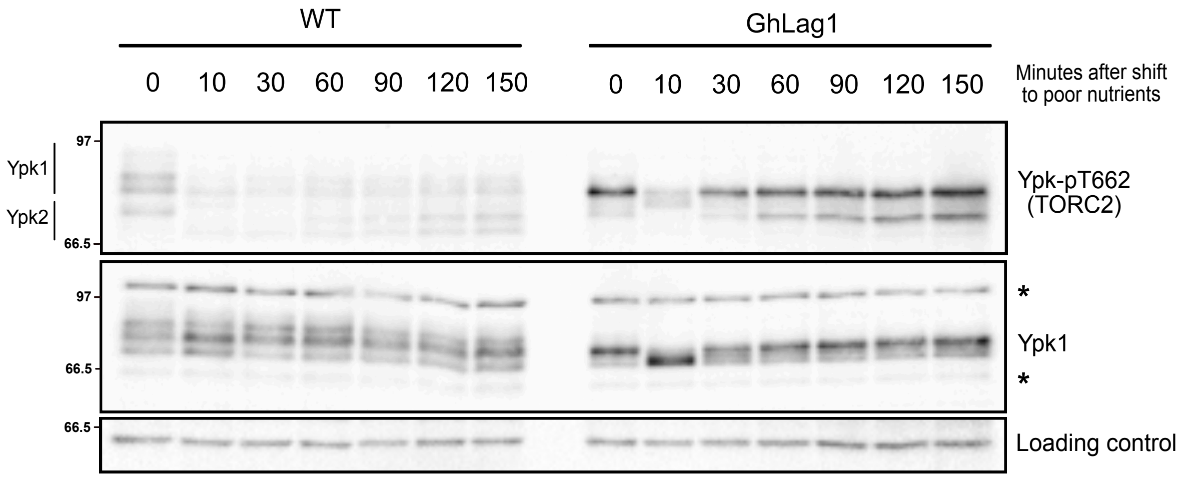
